# Lumbar V3 Interneurons Provide Direct Excitatory Synaptic Input Onto Thoracic Sympathetic Preganglionic Neurons, Linking Locomotor And Autonomic Spinal Systems

**DOI:** 10.1101/2023.04.06.535901

**Authors:** Camila Chacon, Chioma V Nwachukwu, Narjes Shahsavani, Kristine C Cowley, Jeremy W Chopek

## Abstract

Spinal cord injury (SCI) is a life-altering event causing sensation loss, motor paralysis and impaired autonomic function. Electrical spinal cord stimulation has shown promise for restoring lost motor, and impaired autonomic function, despite targeting lumbar locomotor networks. Sympathetic preganglionic neurons (SPNs), primarily located in the intermediate laminae of thoracic and upper lumbar segments (T1-L2), provide neural input to excite sympathetic tissues and organs that provide homeostatic and metabolic support during movement and exercise. We hypothesized that ascending propriospinal neurons located in the lumbar spinal cord provide synaptic input to thoracic SPNs, providing a spinal neural mechanism explaining improved motor and autonomic function in response to spinal cord stimulation. Here, we demonstrate that synaptic contacts from locomotor-related V3 interneurons (INs) are present in all thoracic laminae. Injection of an anterograde tracer into lumbar segments demonstrated that 8-20% of glutamatergic input onto SPNs originated from lumbar V3 INs and displayed a somatotopographical organization of synaptic input to thoracic SPNs, with rostral lumbar V3 INs projecting to rostral thoracic, and caudal lumbar V3 INs projecting to caudal thoracic SPNs. Whole cell patch clamp recording in SPNs demonstrated prolonged depolarizations or action potentials in response to optical activation of either lumbar V3 INs in spinal cord preparations or in response to optical activation of V3 terminals in thoracic slice preparations. This work demonstrates a direct intraspinal connection between lumbar locomotor and thoracic sympathetic networks and suggests communication between motor and autonomic systems may be a general function of the spinal cord.

**Key points:**

- We provide direct anatomical and electrophysiological evidence of an ascending intraspinal synaptic connection between lumbar motor and thoracic sympathetic autonomic neural systems.
- These connections are formed between lumbar locomotor related V3 interneurons and thoracic sympathetic preganglionic neurons (SPNs)
- V3 synaptic input accounts for ∼ 20% of glutamatergic input to SPNs.
- Optical activation of lumbar V3 interneurons elicit action potentials in thoracic SPNs.
- This intraspinal pathway may explain why electrical stimulation of the lumbar region in persons with long-standing motor complete spinal cord injury improves both motor and sympathetic function.
- These findings suggest communication between motor and autonomic systems may be a general feature of spinal cord function and await further research to explore this concept.

## INTRODUCTION

Spinal cord injury (SCI) is a life-altering event, resulting in the loss of sensation and motor paralysis. In addition, persons living with SCI face a lifetime of impaired automatic (autonomic) bodily functions that affect every aspect of daily living. These impairments include reduced temperature and blood pressure (BP) regulation, and inability to activate sympathetic nervous system responses needed during movement and exercise (e.g., increased heart rate (HR), activation of adrenal glands). Spinal electrical stimulation has emerged as a powerful therapeutic to improve motor function after SCI even when applied years after injury. A second critical yet unanticipated set of observations that emerged from electrical stimulation trials targeting lower limb motor centres (lumbar SC), was that it could also improve many autonomic body functions mediated by thoracic spinal sympathetic neurons, including improved blood pressure (BP) and temperature regulation, whole body metabolism and even peak upper body exercise performance (reviewed by (Flett, Garcia et al. 2022)). The underlying neural mechanisms contributing to these functional gains are unknown.

Normally, these autonomic sympathetic preganglionic neurons (SPNs) of the thoracic SC are controlled by autonomic neurons in the rostroventral lateral medulla (RVLM) of the brainstem which has been proposed as a key integration site and descending command centre for cardiovascular control (Loewy and Spyer 1990, Schreihofer and Sved 2011), thermoregulation (Blessing, Yu et al. 1999, Morrison, Sved et al. 1999, Bartness, Shrestha et al. 2010, Bartness, Vaughan et al. 2010) and regulation of lipolysis from white adipose tissues (Bartness, Liu et al. 2014). There is considerable anatomical and electrophysiological evidence supporting the RVLM as a key integration centre for each of these metabolic and homeostatic functions separately and there is also evidence suggesting that the same or overlapping subsets of brainstem and SPNs may provide neural input to multiple tissues simultaneously, including white adipose tissue and muscle or white adipose tissue and the adrenal glands, etc., (Strack, Sawyer et al. 1988, Loewy and Spyer 1990, Smith, Jansen et al. 1998, Kerman, Enquist et al. 2003, Kerman 2008, Llewellyn-Smith and Verberne 2011, Adler, Hollis et al. 2012, Cowley 2018). This region of the RVLM is also the final relay for descending locomotor command signals elicited by stimulation of the MLR (Jordan, Liu et al. 2008). Chemogenetic activation of serotonin neurons in the parapyramidal region elicits hindlimb locomotion and simultaneously, increases BP which precedes and outlasts each bout of locomotion in a genetic decerebrate rat model [(Armstrong, Nazzal et al. 2017) ; cf Fig 4B in (Cowley 2018)].

However, given the absence of descending command signals in ‘complete’ SCI, clinical observations of improved sympathetic autonomic functions suggests that intraspinal, ascending neural connections between lumbar locomotor and thoracic sympathetic neural circuitry become activated during lumbar electrical stimulation. We have previously demonstrated it was possible for completely intraspinal (propriospinal) neurons to relay descending locomotor commands from the brain (Cowley, Zaporozhets et al. 2008, Cowley, Zaporozhets et al. 2010). Further, during fictive locomotion, the lumbar spinal cord is capable of entraining both thoracic (Le Gal, Juvin et al. 2016) and cervical (Juvin, Le Gal et al. 2012) ventral root activity via propriospinal neurons. However, whether propriospinal relays can integrate information between lumbar locomotor-related spinal neurons and sympathetic autonomic spinal neurons has not been investigated and the underlying intraspinal neural mechanisms mediating these clinical improvements are unknown. We recently developed a conceptual framework to better understand the neural mechanisms integrating locomotor and sympathetic autonomic functions (Cowley 2018). We proposed that after SCI, activation of SPNs may be mediated in concert with activation of locomotor-related neurons within spinal locomotor pattern generator (CPG) circuitry as part of an integrated spinal locomotor-sympathetic network. We hypothesized that this coordinated increase in sympathetic output occurs through propriospinal projections to sympathetic nuclei that are activated in parallel with lumbar locomotor-related interneurons [c.f. Fig 4 in Flett et al., (Cowley 2018, Flett, Garcia et al. 2022)].

In order to examine whether an anatomical and functional propriospinal pathway exists to coordinate activity in locomotor-related and autonomic sympathetic neural circuitry, we chose to focus on one of the 11 cardinal classes of genetically identified spinal neurons (Jessell 2000). We focused on the V3 population of spinal neurons because the V3 population of interneurons (INs) are important for stabilizing the frequency of locomotion (Zhang, Narayan et al. 2008), are excitatory, glutamatergic and project both ipsilaterally and contralaterally (Zhang, Narayan et al. 2008, Blacklaws, Deska-Gauthier et al. 2015, Chopek, Nascimento et al. 2018). Computational modeling suggests that V3 INs promote left-right synchrony (i.e., gallop) during high-speed locomotion and have ascending projections from lumbar to cervical regions, providing an anatomical basis for this coordination (Rybak, Dougherty et al. 2015, Danner, Wilshin et al. 2016, Danner, Shevtsova et al. 2017, Danner, Zhang et al. 2019, Zhang, Shevtsova et al. 2022). Here we demonstrate a direct intraspinal neural connection between spinal locomotor related V3 neurons in the lumbar spinal cord and thoracic SPNs. Preliminary results were presented previously in abstract and thesis form (Chacon 2022, Nwachukwu, Chacon et al. 2022).

## METHODS

### Animals

Experimental protocols used complied with the guidelines set by the Canadian Council on Animal Care and were approved by the University of Manitoba animal ethics committee. Anatomical and immunohistochemical observations of V3 soma and nerve terminals, BDA and CTB injections were performed in 9 adult Sim1^Cre/+^;Rosa26^floxstopTdTom/+^ (Sim1TdTom; Zhang et al., 2008) mice of both sexes. Adult female and male Sim1^Cre/+^;Rosa26^floxstopTdTom/+^ mice were crossed with Ai32 mice (Gt(Rosa)26floxstopH134R/EYFP/+, Jackson Laboratory, Stock No. 012569) to generate Sim1^Cre/+^;Rosa26^floxstopTdTom/+^_/_^Gt(Rosa)26floxstopH134R/EYFP/+^ (Sim1TdTom/Ai32 or Sim1TdTom/ChR2). In *vitro* optogenetic electrophysiological experiments were performed in 23 neonatal (P1-P16) Sim1TdTom/ChR2 mice of both sexes.

### BDA and CTB injections

#### Recovery surgery protocol for antero- and retrograde dye injections

Nine adult mice of both sexes were subjected to spinal injection of BDA at spinal segments L2, L3 or L4/L5 for anterograde or CTB-A488 at spinal level T8 for retrograde tracing. All laminectomies and spinal injections were performed aseptically under anaesthesia induced by 5% isoflurane and maintained with 3% isoflurane. Briefly, the skin overlying the vertebrae of interest was opened with a scalpel, back muscles were blunt dissected to reveal the posterior spinous processes and forceps used to perform a single vertebrae laminectomy to expose the spinal cord. Once the dura was opened with a 27-gauge needle, 1 μL of either 1% biotin dextran amine (BDA-10,000 mW, Thermo Fisher Scientific Cat# D1956, RRID: AB_2307337) or cholera toxin B Alexa 488 (CTB-A488, Molecular Probes, Cat # C34775) was injected into the exposed spinal tissue. BDA was injected near the left midline using a stereotaxic micromanipulator and at a depth of 700-900 µm from the dorsal surface of the spinal cord. Injection was delivered using a motorized pico-injector with a 75RN SYR 5 µl Hamilton syringe fitted with a single-barrel borosilicate capillary glass (A-M Systems INC.) pulled to a ∼120 µm tip. The dye was injected over a 5-minute period, the needle remained in place for an additional 10 minutes to prevent dye withdrawal back into the pipette. Each animal received slow-release buprenorphine (0.5 mg/kg), glucose solution (0.02 ml/g), was placed in a heated cage for 2 hours until fully functional and then returned to their home-cage where health was monitored daily. Seven days later, animals were terminally perfused and processed as described in “Immunohistochemistry and Imaging”.

### Immunohistochemistry and Imaging

#### Immunohistochemistry protocol

Under inhalant isoflurane anesthesia, adult Sim1TdTom mice were transcardially perfused with 4% PFA in phosphate-buffered saline (PBS). Spinal cords were dissected free and placed in 4% PFA overnight, and subsequently cryoprotected in 30% sucrose. For sectioning, the thoracic spinal cord was divided into two approximately equivalent-sized portions, with the rostral portion containing ∼T1 – T6/7 and the caudal portion ∼T7 – T13. Lumbar portions were sectioned in one contiguous section at 30 µm. Thoracic spinal cord was sectioned at 18 µm (BDA-injected mice) and 30 µm (all other mice) using a cryostat and mounted on glass slides for immunohistochemistry. To characterize V3 neuron somas throughout thoracic and lumbar segments, images were obtained on a Zeiss Axioscope 40 and every third section (∼ every 450 µm) per slide was examined and/or counted (n = 5 mice). To characterize V3 terminals, SPNs and VGluT2 expression in the thoracic spinal cord, every fifth section (rostro-caudal distance of ∼90 µm between sections) was processed for immunohistochemical visualization of TdTomato and ChAT and VGluT2, with the remainder preserved for other purposes. For BDA-injected mice, every fifth section was examined, representing inter-section distances of ∼ 90 µm. All sections were washed three times for 10 minutes each in 0.2% PBS-T, followed by 10-minute antigen retrieval (55 °C incubation in Tris-EDTA buffer) and an additional three washes in 0.2% PBS-T for 10 minutes each. A 10% normal donkey serum block in 0.2% PBS-T was applied for one hour. Sections were then incubated with primary goat anti-dsRED (1:1000, Takara Bio, Cat# 632496, RRID: AB_10013483), goat anti-ChAT (1:400, Millipore, Cat# AB114P, RRID: AB_2313845) and guinea-pig anti-VGluT2 (1:500, Millipore, Cat# AB2251-I, RRID: AB_2665454) for 72 hours at 4 °C. Sections were then washed three times for 10 minutes each in 0.2% PBS-T followed by incubation with appropriate secondary antibodies conjugated to Alexa 488, Cy3 and Alexa 647 (1:500; Molecular Probes, Cat# A-11055, RRID:AB_2534102; Jackson Immuno Research Labs, Cat# 711-165-152, RRID:AB_2307443; Jackson Immuno Research Labs, Cat# 706-605-148, RRID:AB_2340476, respectively) for two hours. Sections were then washed one time for 10 minutes in 0.2% PBS-T followed by two washes for 10 minutes each in Tris-HCL 50 mM, then dried, mounted in Vectashield hardset mounting medium (Vector Laboratories) and cover slipped. For mice injected with BDA, sections were processed as described but immunolabelling for VGluT2 was removed and the secondary antibody Streptavidin647 (1:500, Thermo Fisher Scientific, Cat# S-21374, RRID: AB_2336066) was used at the appropriate step to label for BDA. Similarly, in a subset of mice that received thoracic injections of CTB, lumbar spinal cords were washed, mounted in Vectashield hardset mounting medium and cover slipped.

#### Image Analysis

Sections confirming BDA and CTB injection sites and CTB positive lumbar V3 INs were imaged using a Zeiss Axio Imager Z.2 upright microscope. BDA injection sites were reconstructed using tiled images obtained under 10x magnification within Zen Blue software (Version 3.5). Images were used to confirm injection site and determine rostro-caudal and transverse distribution of injected dye. For CTB-injected mice, CTB-A488^+^/tdTomato^+^ somas in lumbar cord were counted by creating tiled, Z-stack images (7 – 10 z-stack images through a focusing range of 6 – 9 µm) at 10x magnification and counted in ZenLite software (Version 3.6). For each animal, 9 – 10 sections were imaged and analyzed throughout the entire lumbar spinal cord per animal (L1 – L6; n=4 animals). To correspond with previous anatomical observations of three distinct V3 sub-populations (Zhang, Narayan et al. 2008, Borowska, Jones et al. 2013), tiled images of each transverse spinal cord section were divided into dorsal, intermediate and ventral regions for both sides of the lumbar spinal cord to compare double labelled somas ipsi- and contra-lateral to the injection site. For a neuron to be classified as a V3 IN labelled with CTB, either a distinguishable overlap of the neuron’s soma with CTB fluorescence was present or the neuronal soma had to have perinuclear labelling with 4-5 distinct vesicle speckles within demarcations of the soma (Conte, Kamishina et al. 2009).

A Zeiss Airyscan LSM 880 confocal microscope was used to image VGluT2/TdTom^+^ or BDA/TdTom^+^ neuronal processes apposing SPNs (ChAT^+^ neurons) in thoracic IML. For each rostral (∼T1 – T6/7) and caudal (∼T6/7 – T13) region, 14 – 20 ROIs containing the IML, with at least 90 µm of inter-section distance, were selected to obtain a representative rostro-caudal distribution of IML regions and SPNs per mouse (n=4). Each IML area was imaged [7 – 14 Z-stack images through a focusing range of 3.6 – 12.8 µm) at 63x magnification. Laser settings were optimized and replicated within each animal, for consistency between images. Left and right IML SPNs were selected and imaged based on visibility of at least one SPN soma per ROI. Z-stack images were then imported into IMARIS software for 3D reconstruction and quantitative analysis using the “Surface” tool. Synaptic boutons (VGluT2) less than 1 µm from SPN somas were quantified and considered doubled labelled if there was overlap with TdTom^+^ surfaces. Similarly, doubled labelled contacts of BDA/TdTom^+^ were considered as synaptic boutons when less than 1 µm from SPN soma. To examine presumed V3 IN terminal density in thoracic IML, one half of thoracic grey matter was tile-imaged using a 20x objective lens on the Zeiss Airyscan LSM 880 confocal microscope (2 – 3 Z-stack images through a focusing range of 2.49 – 4.99 µm) per thoracic section (∼12 images, taken at regularly spaced intervals, corresponded with each thoracic spinal region, n = 3 mice). Images were then reconstructed and analyzed in IMARIS. “Spots” detection method was used to filter and select presumed V3 IN terminals. Cell bodies and long neuron processes were excluded from quantification in IMARIS using its size and shape exclusion and re-iterative learning features. A region of interest set in IMARIS around the IML (278 x 277 µm) was used to analyze and quantify V3 neuron synaptic terminal density within each spinal tissue section. Individual X–Y coordinates for each synaptic terminal were exported to Excel and GraphPad Prism for subsequent analysis. XY coordinates for each synaptic terminal were also exported and transformed into contour plots using RAWGraphs (https:app.rawgraphs.io).

### Electrophysiology

#### Spinal cord slice preparations

Experiments were performed on spinal sections isolated from 23 neonatal (P1-P16) Sim1^Cre/+^;Rosa26^floxstopTdTom/+^/Gt(Rosa)26^floxstopH134R/EYFP/+^ (referred to as Sim1TdTom/ChR2) mice. Isolation and preparation of sections, and electrophysiological recording methods have been previously described (Chopek, Nascimento et al. 2018). Briefly, animals were anaesthetized with isoflurane, decapitated at the medulla-spinal cord junction and spinal cords dissected out in ice-cold dissecting artificial cerebrospinal fluid (aCSF), composed of (mM): KCl (3.5), NaHCO_3_ (35), KH_2_PO_4_ (1.2), MgSO_4_ (1.3), CaCl_2_ (1.2), glucose (10), sucrose (212.5), MgCl_2_ (2.2), and equilibrated to pH 7.4 with 95% 0_2_ & 5% C0_2_. Once dissected free, thoracic spinal cords were immediately secured in agarose and sectioned at 350 μm using a vibratome (Leica VT1200S, Leica) and then incubated in warm (30 °C) aCSF for a minimum of 30 minutes before performing electrophysiological experiments. Incubation and recording aCSF was composed of (mM): NaCl (111.0), KCl (3.085), D-glucose (10.99), NaHCO_3_ (25.0), MgSO_4._7H_2_O (0.31), CaCl_2_ (2.52), KH_2_PO_4_ (1.1), equilibrated to pH 7.4 with 95% O_2_ & 5% CO_2_.

#### Intact spinal cord preparations

Spinal cords were dissected free from Sim1TdTom/ChR2 mice as described above in ‘Slice preparations’ but remained longitudinally intact with dorsal and ventral roots attached. Once free, connective tissue was carefully removed from spinal cord tissue and secured ventral side up with fine insect pins. To retrogradely label SPNs, rhodamine dextran amine dye (RDA) was applied to cut T6-T8 ventral roots on one or both sides of the spinal cord using glass suction pipettes with internal diameters of 100-120 µm. Ventral roots were cut close to their exit from the spinal cord to minimize labeling time. Retrograde labeling of thoracic SPNs continued in the dark at room temperature for at least 3 hours (Szokol, Glover et al. 2008). A block of agar was prepared in advance with one side of the block cut with a scalpel to provide a 30-degree incline. The spinal cord was then mounted on the agar block and fixed in place with acrylic glue to expose the corresponding labeled thoracic spinal segment. The spinal cord at the level of the agar block was then sectioned with a vibratome. This portion of the spinal cord was then glued to a sylgard-coated (Sylgard, Dow Corning, MI, USA) recording chamber designed and 3D printed in-house specifically for these experiments. In particular, the obliquely cut surface of the spinal cord was placed on a sylgard ‘ramp’ to enable visualisation and patch-clamp recordings of SPNs under fluorescence while preserving and maintaining continuity with the lumbar spinal region for optical stimulation.

#### Whole cell patch-clamp recordings and optogenetic stimulation

Slices or spinal cord preparations were transferred to a recording chamber mounted on a Zeiss AxioExaminer microscope and perfused with oxygenated room-temperature aCSF. Cells were visualized using a 20x wide aperture (1.2 nA) water-immersion objective lens, a CCD camera (CoolSNAP EX OCD Camera, Photometrics, AZ) and Slidebook 6.0 software (Intelligent Imaging Innovations, CO, USA, RRID:SCR_014300). Patch pipettes were pulled with a P-97 Sutter puller and those with 4-6 MΩ resistances were filled with the following (mM): K-gluconate (128); NaCl (4); CaCl_2_ (0.0001); Hepes (10), glucose (1); Mg-ATP (5); and GTP-Li (0.3). Whole cell patch-clamp recordings were made under current-clamp using a Multiclamp 700B amplifier (Molecular Devices, California, USA, RRID: SCR_014300). Recordings were low pass filtered at10 kHz and acquired at 25 kHz with CED Power 1401 AD board and displayed and recorded using Signal software (Cambridge Electronic Design, Cambridge UK). Before performing an optical stimulation protocol, rheobase (defined as the minimum current to elicit a single AP was collected to determine cell excitability). Before the optical stimulation protocol, we recorded rheobase and repetitive firing in response to 1 second depolarizing current steps from each SPN. In slice preparations, presumed SPNs (small neurons visualized in the IML, with TdTom fluorescence noted near the soma) were patched and responses recorded. Using a spatial light modulator system as previously described (Chopek, Nascimento et al. 2018, Chopek, Zhang et al. 2021), a region of interest slightly greater than the SPN soma was created for optical stimulation of presumed V3 terminals apposing the patched SPN. Blue light was delivered in five 500 ms pulses at 5 Hz at a laser power of ∼2.5 mW. For intact cord preparations with thoracic surface exposed, SPNs that were retrogradely labelled with RDA were patched and responses recorded. A region of interest covering the ventral L2 segment was created for optical stimulation of presumed ventral L2 V3 neurons and likely axons of passage from caudal lumbar segments. Blue light was delivered in five 500 ms pulses at 5 Hz at a laser power of ∼2.5 mW. In a subset of slice experiments, lumbar V3 neurons were patched and optically activated to confirm that optical stimulation resulted in AP generation in V3 neurons.

### Statistics

#### Statistical Analysis

Data are presented as means ± SD. As is common for discovery experiments, no statistical method to predetermine sample size and no randomization or blinding procedures were used. Statistical comparisons using t-tests or one-way ANOVA and Tukey’s multiple comparisons tests were performed using GraphPad Prism (V9.5 for MacOS, GraphPad Software, San Diego, California USA, www.graphpad.com). Tests for normality were used to select either parametric or non-parametric tests. Statistical significance was set to p < 0.05.

## RESULTS

### V3 IN cell bodies located throughout the thoracic spinal cord

V3 INs cell bodies have been localized within caudal thoracic and upper lumbar spinal regions (Zhang, Narayan et al. 2008, Borowska, Jones et al. 2013) and electrophysiological evidence suggests a role for V3 INs in lumbar locomotor activity (reviewed by (Ziskind-Conhaim and Hochman 2017), but their presence or function(s) in more rostral thoracic regions has not been investigated. Thus, we examined the anatomical distribution of V3 cell bodies throughout the thoracic and lumbar spinal cord (Figure 1). Our findings regarding V3 soma in lumbar regions were consistent with previous reports, with three subpopulations (dorsal, ventral and intermediate subpopulations) in the rostral lumbar and two populations (intermediate and ventral) observed in caudal lumbar regions (e.g., Fig. 1Cg vs. 1Ch).

**Figure 1.**
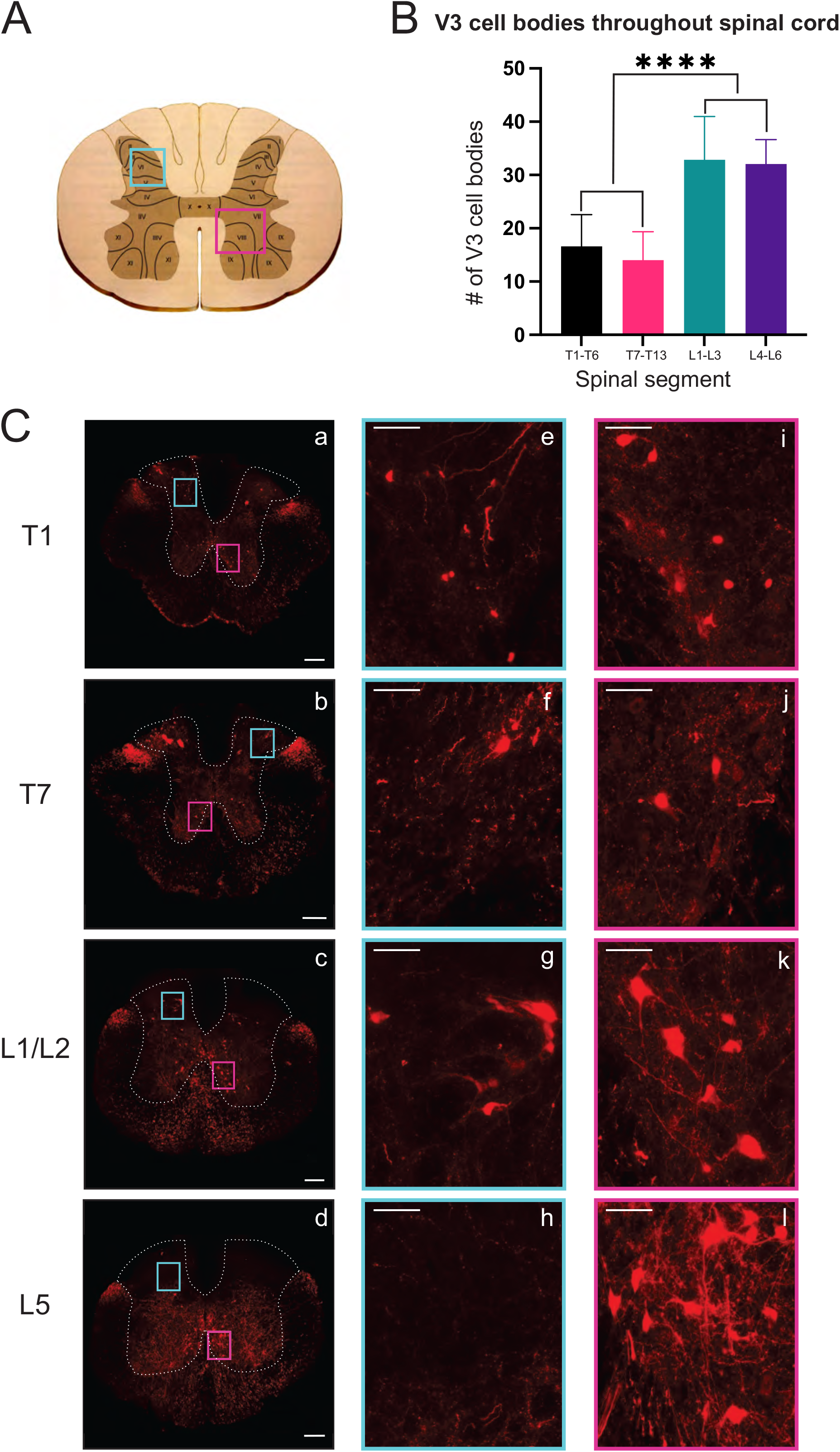
V3 neuronal cell bodies distributed throughout the thoracic and lumbar spinal regions. A) Schematic and cartoon demonstrating Rexed’s Laminae and regions from dorsal (turquoise boxes) and ventral (fuschia boxes) regions for images shown in C. B) Summary data showing mean numbers (± SD) of V3 cell bodies in rostral and caudal thoracic and lumbar regions. Similar numbers of V3 cell bodies were observed within rostral and caudal thoracic and lumbar regions but differed between thoracic and lumbar regions (P <0. 0001), C) V3 cell bodies were observed at each spinal level in ventral regions (1C, i-l) and in dorsal regions at T1 (1Ce), T7 (1Cf) and L1/2 (1Cg) but were absent at L5 (1Ch). Scale bar in a-d = 200 μm, and = 50 μm for remaining images.

In the thoracic spinal cord, we observed V3 soma throughout T1-T13 at similar dorsal and ventral transverse locations as observed in rostral lumbar regions (Fig. 1). V3 soma were not observed within thoracic IML regions. V3 soma in the thoracic region were smaller and less densely distributed compared to V3 cell bodies located in the lumbar region (Compare panels for T1, T7, L1/2 and L5 in Fig. 1C – and graph in 1B). On average, similar numbers of V3 cell bodies were observed per 30 µm section in rostral versus caudal thoracic segments (16.6 ± 5.94 V3 soma in T1-T6; 14.0 ± 5.36 V3 in T7-T13; Fig. 1B & Fig. 1C T1vs Fig. 1C T7, n=4). The highest number of V3 somas were observed in lumbar sections (i.e., L1 – L3 with 32.85 ± 8.14 and L4 – L6 with 32.05 ± 4.59 V3 neurons per section, *p* < 0.0001; Fig. 1B & Fig. 1C L1/2 vs Fig.1C L5). In addition to V3 cell bodies, V3 TdTom^+^ fibres were observed throughout the thoracic spinal cord and within IML regions.

We therefore characterized the distribution of presumed V3 synaptic terminals within the rostro-causal thoracic grey matter generally and with a particular focus and analysis of terminals within the IML. To do so we identified TdTom^+^ presumed nerve terminals (1 section located at ∼ 900 µm distance intervals) throughout the thoracic spinal cord, including a pre-defined IML region in Rexed lamina VII (12 sections per animal, n = 3). V3 terminals were identified at all rostro-caudal levels of the thoracic region, within the dorsal and ventral horns, with an expected high density in lamina IX, close to the somatic motor nuclei (e.g., ∼T10 in Fig. 2B). For consistency, a pre-defined area (278 μm x 277 µm) of the IML was selected for quantifying the number of V3 terminals for each section (Schematic in Fig. 2A, blue box). Each TdTom^+^ terminal was marked with an XY coordinate and exported from IMARIS to develop contour plots (heat maps) of the relative distribution of terminals throughout grey matter (see representative example of Fig.2A transformed to terminal only map in Fig.2B). Numbers of terminals within the grey matter ranged from ∼2500 – ∼17, 700 (mean = 9646, 7577, 5526 for mouse 1, 2 and 3 respectively) and within the IML, terminal numbers ranged from ∼500 – ∼2400 per section (mean= 1279, 1166, 875 for mouse 1, 2 and 3 respectively), depending on the rostrocaudal level of the section. In particular, there were higher numbers of terminals in the pre-defined IML region in rostral (∼ ≥ 1,000) compared to caudal (∼ ≤ 1,000) thoracic SC (Compare heat maps of T1 to T12/13 in Fig. 2B and numbers in Fig. 2C) with highest numbers observed between ∼T4 and T6 (Fig. 2C).

**Figure 2.**
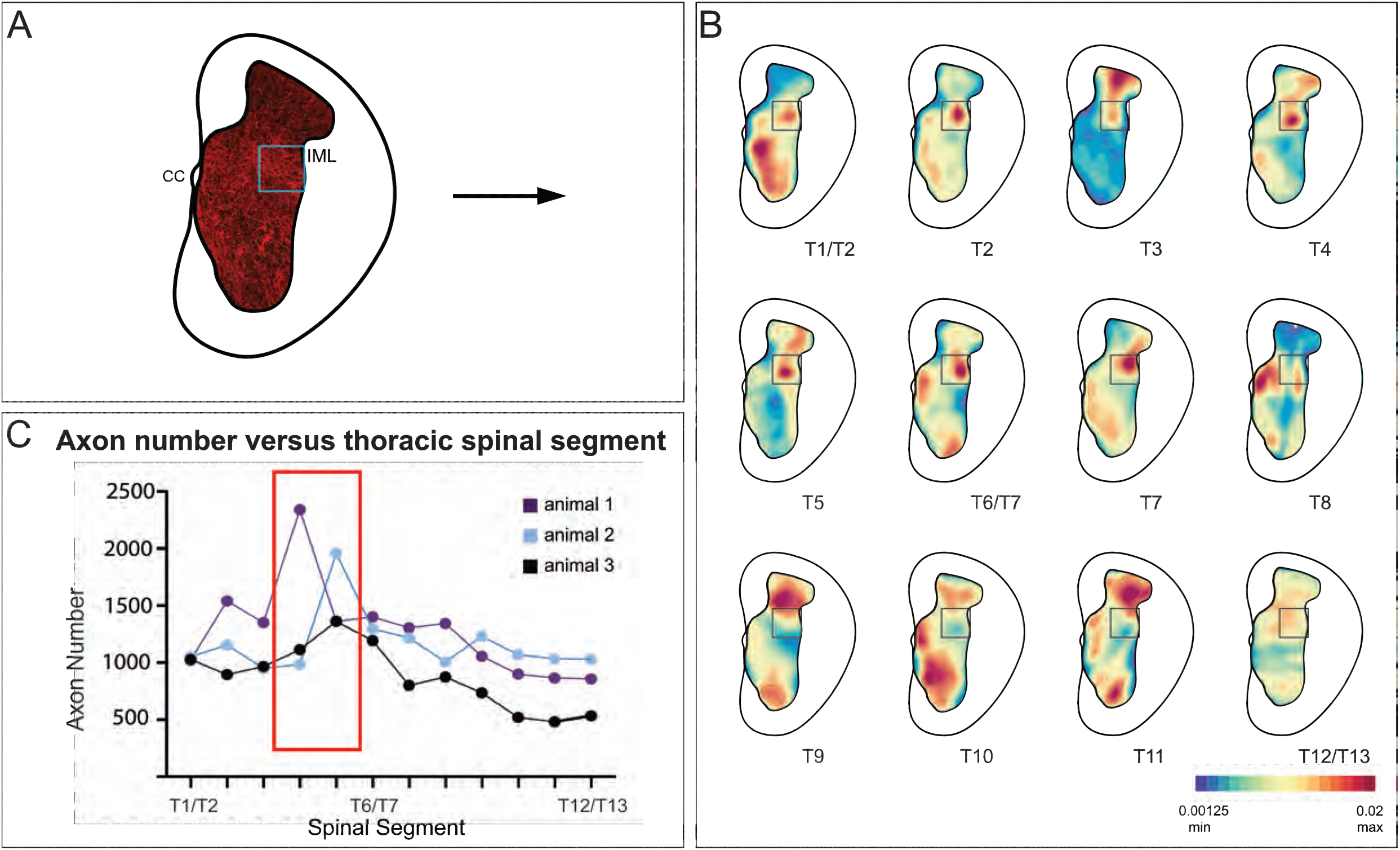
V3 neuron terminals observed within the IML at all thoracic spinal segments, with highest density observed at mid-thoracic levels. A) Schematic demonstrating V3 cell bodies and terminals observed in grey matter of spinal cord. XY coordinates of V3 terminals were exported to create contour maps in IMARIS. B) Representative contour plots of relative V3 terminal densities within grey matter at each thoracic spinal level from one adult mouse. Note the high density of V3 terminals in the IML and throughout grey matter. C) Graph summarizing numbers of V3 terminals observed in IML ROI (278 μm x 277 µm) at each thoracic spinal level for each animal examined (n=3 animals, 12 sections per animal). Numbers of terminals within the grey matter ranged from ∼2500 – ∼17, 700, and within the IML, terminal numbers ranged from (∼500 – ∼2400) per section, depending on the rostrocaudal level of the section.

Together these results demonstrate that V3 cell bodies are located throughout the thoracic spinal cord, distributed in similar anatomical populations, but at approximately half the size and number than observed in rostral lumbar regions. These results also demonstrate that there is a large proportion of V3 neuronal terminals within the IML region of the thoracic spinal cord (presumed V3 IML puncta account for 14.3%, 15.9% and 21.2% of all puncta within grey matter of one side of spinal cord for animal 1, 2 and 3 respectively).

### 20% of glutamatergic input apposing thoracic SPNs arises from V3 interneurons

In neurologically intact preparations, VGluT2^+^ terminals constitute the main, if not only, glutamatergic nerve terminal input onto SPNs (Llewellyn-Smith, Phend et al. 1992, Llewellyn-Smith, Martin et al. 2007). In order to determine whether any excitatory V3 terminals observed within thoracic IML directly apposed SPNs, we examined our sections labelled with ChAT (SPNs), VGluT2 and DsRed. Confocal z-stack images from rostral (Figure 3A) and caudal thoracic regions (Figure 3C) were imported into IMARIS software for 3D reconstruction. Using the 3D reconstructions, we calculated and plotted the number of VGluT2^+^ and double-labelled VGluT2^+^/DsRed^+^ synaptic terminals apposing SPNs for sections from rostral and caudal thoracic regions for each animal (Fig. 3E). A total of 56-80 SPN sections were reconstructed and analyzed for each rostral and caudal thoracic region per mouse (n=4). On average, 107.2 VGluT2^+^ contacts apposed each SPN (range: 0 – 645 VGluT2^+^ contacts). Over all sections examined for each animal, 16.9% – 26.6% (n=4 mice) of VGluT2^+^ synaptic terminals were double labelled with DsRed, indicating that these terminals arose from V3 neurons (Fig. 3A & C). Similar proportions of excitatory V3 synaptic terminals were observed apposing SPNs within rostral versus caudal thoracic regions in each animal (Fig. 3E; *p* = 0.43, n =4).

**Figure 3.**
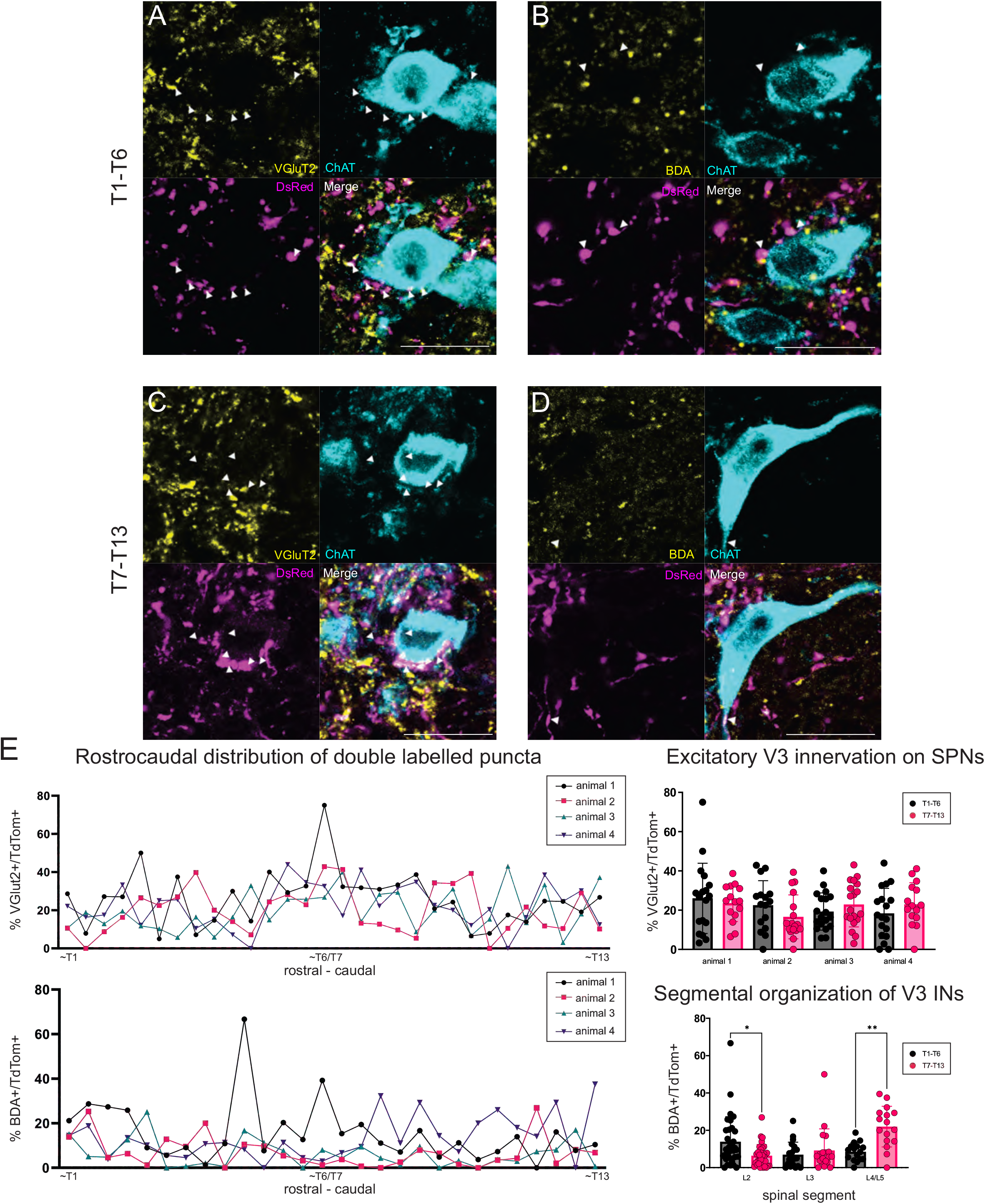
Injection of BDA neuronal tracer into lumbar spinal cord reveals ascending synaptic input from V3 neurons located in the lumbar region. A) Double labelled VGluT2 and TdTom terminals apposing SPNs in rostral thoracic spinal cord. Double labelled neurons indicated with arrowheads. B) Double labelled BDA and TdTom terminals apposing SPNs in rostral thoracic spinal cord. Double labelled neurons indicated with arrowheads. C) Double labelled VGluT2 and TdTom terminals apposing SPNs in caudal thoracic spinal cord. Double labelled neurons indicated with arrowheads. D) Double labelled BDA and TdTom terminals apposing SPNs in caudal thoracic spinal cord. Double labelled neurons indicated with arrowheads. **Scale bar** = 20 μm for panels (A-D). E) Rostrocaudal distribution of percentage of double labelled VGluT2^+^/TdTom^+^ and BDA^+^/TdTom^+^ puncta within thoracic spinal cord. Glutamatergic excitatory V3 IN innervation of SPNs averaged ∼20% per section examined, with no significant difference between T1-T6 and T7-T13 regions (p = 0.43, Brown-Forsythe test, n = 4 mice). In animals with BDA injection at L2, higher percentage of BDA^+^/TdTom^+^ terminals were present in sections from rostral thoracic compared to caudal thoracic regions (* p < 0.0265, n = 2 mice). In contrast, BDA injections in L4/5 demonstrated higher percentages of BDA^+^/TdTom^+^ terminals in sections from caudal thoracic compared to rostral thoracic regions (** p < 0.0019). L3 BDA injections showed no significant difference in percentage of BDA^+^/TdTom^+^ terminals in sections from either rostral or caudal thoracic regions (p < 0.977).

We noted that SPN size varied throughout the thoracic spinal cord, therefore we analyzed the number of V3 terminals apposing each SPN, normalized to observed SPN surface area, and found a similar ratio between rostral (0.63 + 0.52) and caudal (0.64 + 0.63) thoracic SPNs suggesting similar densities of V3 terminals apposing thoracic SPNs (*p* = 0.5) regardless of rostrocaudal location or when normalized for relative size of SPN cell body.

In order to determine if any of the V3 synaptic terminals apposing thoracic SPNs originated from lumbar V3 neurons, in the next series of experiments we injected BDA into lumbar spinal cord and then quantified the proportion of V3 contacts (DsRed^+^) that were also BDA^+^ and apposing ChAT^+^ SPNs. Four animals contributed to this series (2 injected at L2, 1 at L3 and 1 at L4/L5). A typical example is shown in Figure 3 with images taken from the rostral (Fig. 3B) and caudal (Fig. 3D) thoracic regions demonstrating that a portion of V3 synaptic input onto SPNs originates from V3 neurons in the lumbar region (animal injected at L2). Overall, for each section examined, 3.8 – 21.9% of total V3 contacts (DsRed^+^) were also BDA^+^. Demonstrating that lumbar V3 neurons provide ascending synaptic input onto thoracic SPNs, and double-labelled terminals were observed when injected at either L2, L3 or L4/5 (Fig. 3E, bottom panel). Overall, 9.7 % (± 5.1%) of V3 neurons were also labelled with BDA for the entire rostral thoracic region (T1–T6; n=4; range = 5.8 to 17.8%) and 9.8% (± 7.8%) for the caudal thoracic region (T7-T13; n=4; range = 3.8 to 21.9%). Thus approximately half of all V3 contacts on SPNs arose from V3 neurons in lumbar SC.

In this series we also compared the proportion of BDA^+^ and DsRed^+^ presumed terminals in rostral versus caudal thoracic regions to determine if there was any evidence to suggest a somatotopic organization to the ascending input onto thoracic SPNs from lumbar V3 neurons. Thus, we compared the proportion of double-labelled terminals observed in rostral (T1-T6; Fig. 3B) versus caudal (T7-13; Fig. 3D) thoracic regions for BDA injections at rostral lumbar (L2, Fig. 3E) versus caudal lumbar (L4/5; Fig. 3E) sites. Figure 3E demonstrates that there was a greater proportion of BDA^+^ and DsRed^+^ terminals in sections from rostral thoracic versus caudal thoracic regions for animals injected with BDA at rostral lumbar sites (13.9% vs 6.4% respectively) and see also animal injected at L2 in Fig. 3B & 3D, * p < 0.0265, n=2 mice). Conversely, animals injected with BDA at caudal lumbar sites demonstrated a greater proportion of BDA^+^ and DsRed^+^ terminals in sections from caudal compared to rostral thoracic regions (22.0% vs 8.5% respectively, Fig. 3E, n = 1 mouse, ** p < 0.0019). In contrast, a similar proportion of BDA^+^ and DsRed^+^ terminals were observed in sections from rostral and caudal regions in animals with BDA injection at a mid-lumbar (L3) level (6.95 % vs 9.23% respectively, Fig. 3E, n = 1 mouse, p < 0.977). These findings suggest that ascending input from lumbar V3 neurons is somatotopically arranged, as opposed to a pattern of ascending input throughout the entire thoracic region, regardless of V3 lumbar segmental location.

We next aimed to identify which V3 subpopulation(s) innervate thoracic SPNs. To do so, we unilaterally injected CTB targeted to the IML at the T8 segment and counted CTB^+^ V3 INs throughout the lumbar spinal cord (Fig. 4, n = 3). Due to the difficulty in precisely targeting SPNs, which are located near white matter that may also contain V3 axons of passage, we examined each injection site for potential spread and present descriptive evidence from all injections and compared these findings to the one confirmed successful unilateral injection (Fig. 4B) in which no CTB was evident in the white matter. We found widespread CTB uptake in the dorsal, intermediate and ventral V3 subpopulations both ipsilateral and contralateral to each injection site (Fig. 4C & D). Similar percentages of labelled V3 INs were observed in ipsilateral dorsal (73.5% and 73.8%), intermediate (62.7% and 53.3%) and ventral (59.7% and 58.0%) lumbar regions when the confirmed ipsilateral grey matter CTB injection was compared to the three other more widespread CTB injections. Similar percentages of labelled V3 INs were observed in contralateral dorsal (64.3% and 70.0%), intermediate (55.1% and 53.5%) and ventral (67.4% and 66.9%) lumbar regions CTB (Fig. 4E). This suggests that both contralateral and ipsilateral, and dorsal and ventral lumbar V3 INs project near thoracic IML, consistent with previous findings of both ipsilateral and contralateral projections arising from V3 INs to motor related regions in cervical SC (Zhang, Narayan et al. 2008, Borowska, Jones et al. 2013, Blacklaws, Deska-Gauthier et al. 2015, Chopek, Nascimento et al. 2018).

**Figure 4.**
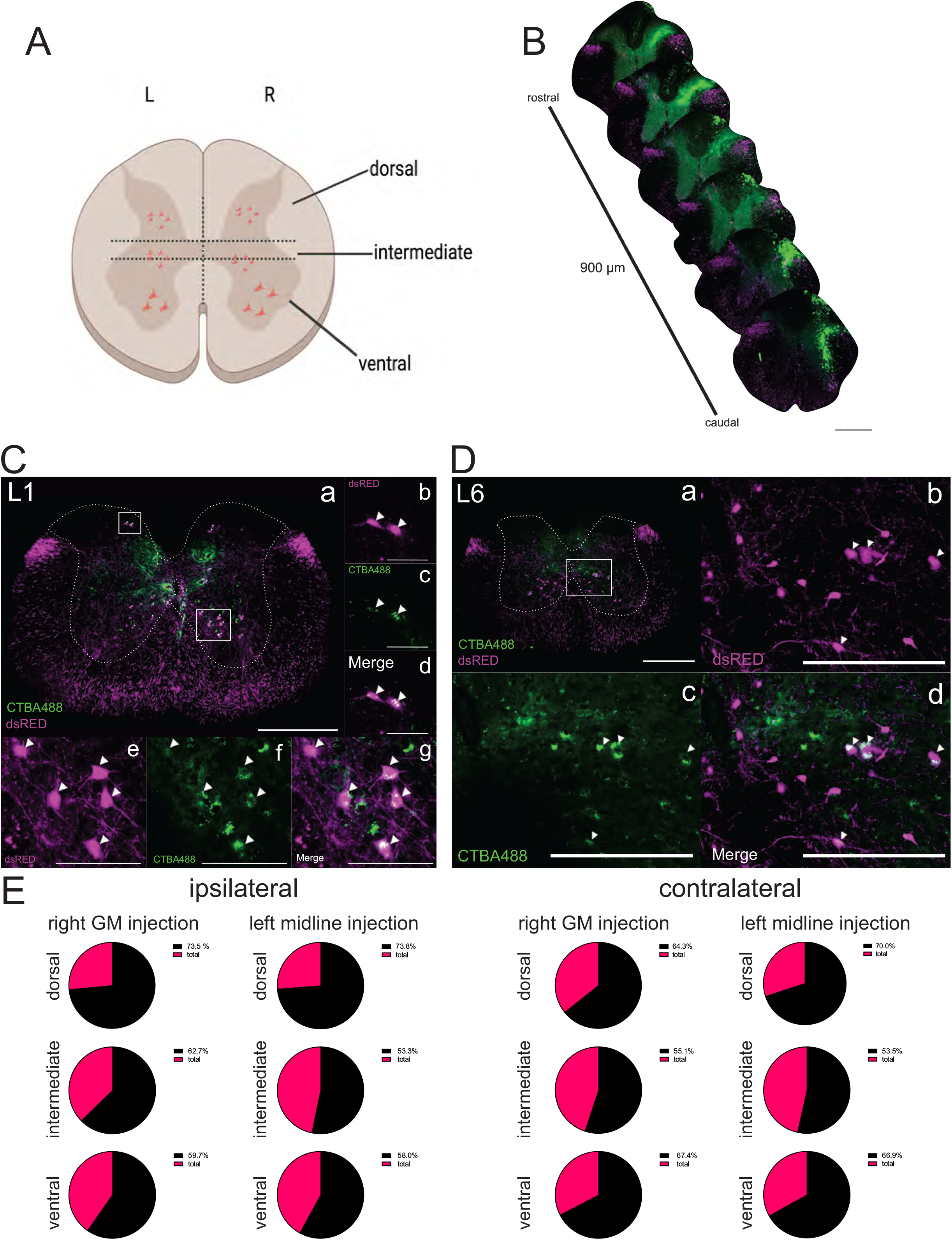
Injection of CTB neuronal tracer targeting IML at thoracic segment T8 suggests all lumbar V3 IN subpopulations project to thoracic spinal cord. A) Schematic of lumbar spinal cord section showing dorsal, intermediate and ventral V3 IN locations. Sections were divided into dorsal, intermediate and ventral regions on left (L) and right (R) sides of the spinal cord and labelled V3 INs were counted based on these regions of interest. B) Reconstructed T8 injection site. CTBA488 (green) was injected unilaterally into the spinal cord. This injection was confined to one side of the spinal cord. **Scale bar** = 500 μm C) Image from L1 segment of spinal cord (Fig. 4Ca) from same animal as in 4B. Enlarged image showing area in box placed over dorsal grey matter demonstrates double labelling of CTBA488^+^ and dsRED^+^ in dorsal V3 INs (4Cb-d for dorsal ROI) as well as in ventral regions (see box over ventral grey matter and corresponding enlarged images in 4Ce-g). **Scale bar** = 125 μm. D) L6 section of lumbar spinal cord (a) from same animal as in 4B. **Scale bar** = 500 μm. Enlarged images (4Db-d) show area outlined by box in ventral grey in 4D demonstrates double labelling of CTBA488^+^ and dsRED^+^, **Scale bar** = 125 μm. E) Summary charts indicate similar proportions of ipsi- and contralateral labelling from injections at the midline (n = 3 injections had bilateral dye spread) versus the animal with injections restricted to one side of the SC (GM, n = 1 injection). Similar proportions of double labeling of CTBA488^+^/dsRED^+^ within V3 INs were also observed in dorsal, intermediate and ventral grey matter ipsilateral and contralateral to injection sites throughout the lumbar spinal cord (n = 9 – 10 sections per mouse).

### Spinal V2a and V0 neurons do not appose SPNs

As part of our investigations to identify potential sources of intraspinal locomotor-related neuronal input onto sympathetic-related neural circuitry, we performed anatomical experiments to determine whether two other genetically defined candidate spinal neuron populations provide synaptic input to thoracic SPNs or the IML region generally. The first population were the locally projecting, glutamatergic V2a neurons involved in left-right coordination and defined by the transcription factor Chx10 (Al-Mosawie, Wilson et al. 2007, Lundfald, Restrepo et al. 2007, Crone, Quinlan et al. 2008, Crone, Zhong et al. 2009). Examined in 2 Chx10:eGFP mice, we observed VGluT2^+^/GFP^+^ fibres within the IML but did not observe any double-labelled (VGluT2^+^/GFP^+^) terminal apposing thoracic SPNs. In these same animals, we did observe VGluT2^+^/GFP^+^ terminals apposing lumbar motoneurons, suggesting that our negative findings regarding thoracic SPNs was not due to a weak fluorescent signal or lack of GFP expression in terminals of Chx10 mice.

In another series of experiments, we examined whether V0 population neurons provide synaptic input onto SPNs. V0 neurons can be subdivided into 3 subpopulations (V0_c_, V0_D_, V0_v_) each with a different neurotransmitter content, projection patterns and roles in locomotion (Moran-Rivard, Kagawa et al. 2001, Pierani, Moran-Rivard et al. 2001, Lanuza, Gosgnach et al. 2004, Zagoraiou, Akay et al. 2009, Talpalar, Bouvier et al. 2013). Thus, in 2 Dbx1Cre:tdTomato mice, in which all three populations are visualized with red fluorescent protein, we did not observe any TdTom^+^ terminals apposing thoracic SPNs. Similar to our observations for the V2a population noted above, we did observe terminals apposing lumbar motoneurons, suggesting that our negative findings regarding thoracic SPNs for the V0 population were not due to a weak fluorescent signal or lack of TdTomato expression in terminals of Dbx1 mice.

### Lumbar and thoracic V3 neurons provide functional excitation to thoracic SPNs

Since our anatomical investigations demonstrated that V3 spinal neurons provide direct synaptic input onto thoracic SPNs, we performed a series of electrophysiological experiments to determine if activation of V3 neurons provide functional excitation of SPNs. To do so, we bred mice to obtain channel rhodopsin expression in V3 INs and used immunohistochemical microscopic examination to confirm channel rhodopsin expression in V3 neurons in Sim1^Cre/+^;Rosa26^floxstopTdTom/+^/Gt(Rosa)26^floxstopH134R/EYFP/+^ mice (Fig. 5A) and that TdTom^+^/Chr2^+^ terminals were present in thoracic IML regions (Fig 5B). We then confirmed that we could elicit action potentials (APs) in lumbar V3 neurons in response to optical stimulation (Fig. 5C, n=10). In 10 out of 10 V3 neurons tested, optical stimulation over the soma of the patched V3 in lumbar SC elicited repetitive firing of the neuron.

**Figure 5.**
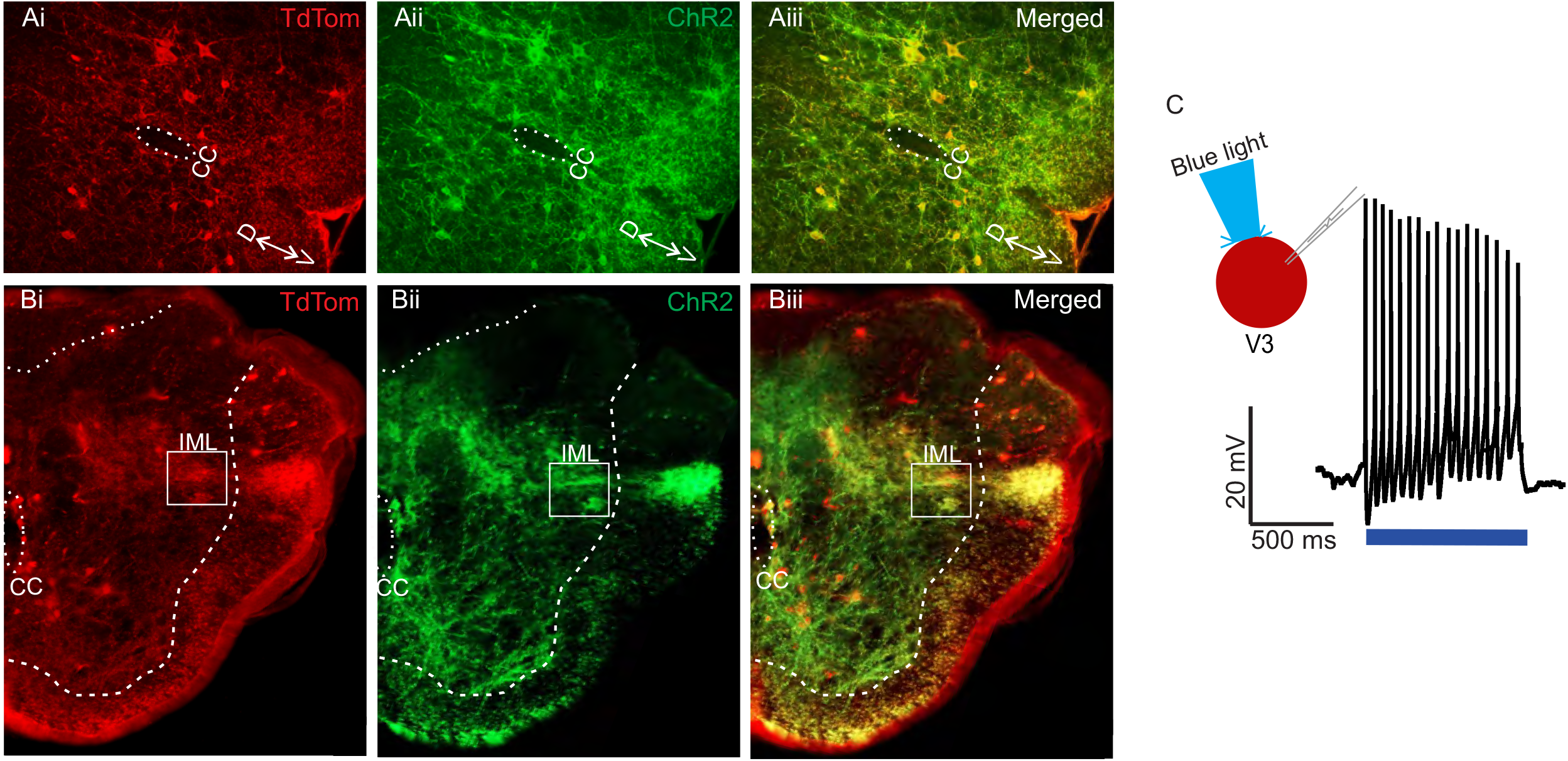
Channel rhodopsin expression in lumbar V3 neurons and nerve terminals in the IML. A) Transverse sections from L2 spinal cord demonstrating V3 neurons (TdTomato Ai, red) also express channel rhodopsin (Aii, green), with merged image in Aiii, yellow. Dorsal (D) and ventral (V) and central canal (CC) shown. B) Transverse sections of T6 spinal cord demonstrate expression of Tdtomato (Bi) in V3 fibres within IML (white box), channel rhodopsin (Bii), and both in merged images (Biii). C) Optical stimulation evokes multiple action potential in whole-cell patch clamped V3 neurons located in thoracic SC. Example from n= 10 cells from 6 mice of either sex.

Next, we determined whether optical stimulation of V3 neuron terminals in thoracic slice preparations could excite putative SPNs in the IML. Examined in 10 presumed SPNs, optical stimulation (480 nm wavelength) of V3 terminals directly over the patched presumed SPN using a focal light source from a spatial light modulator elicited excitatory post synaptic potentials (EPSPs, Fig. 6A n=2), action potentials (Fig. 6B n=4) or no response (n=4, not shown). Presumed SPNs in slice recording had an average rheobase value of 20 ± 5 pA.

**Figure 6.**
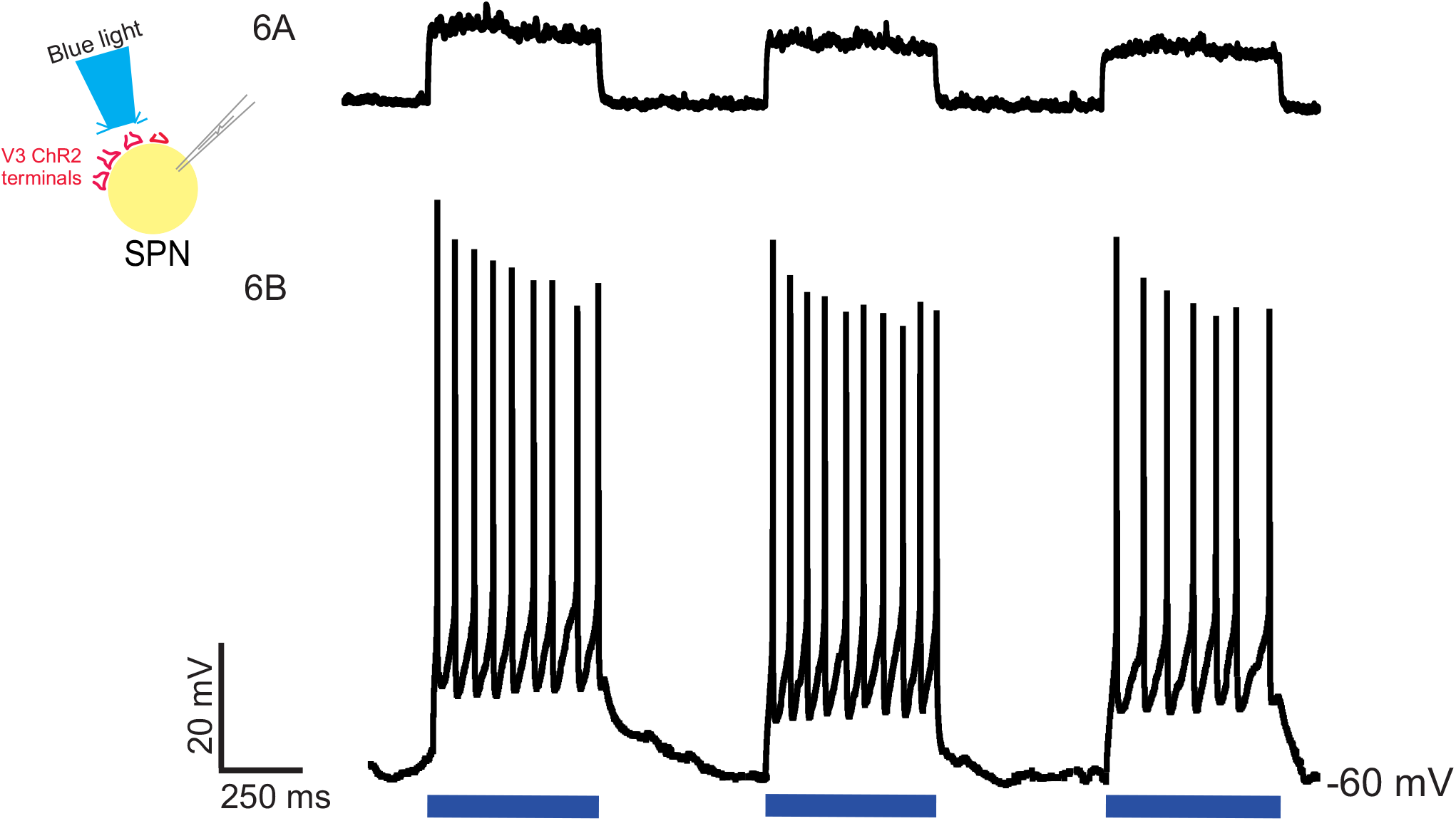
Optical stimulation of V3 nerve terminals elicits excitatory post synaptic responses in thoracic SPNs. Optical stimulation of V3 nerve terminals could elicit EPSPs (A, n=2) or action potentials (B, n=4) in whole cell patched clamped thoracic SPNs. Examples from n= 10 cells from 4 mice of both sexes.

In our final electrophysiological series, we determined whether optical stimulation of lumbar V3 neurons could excite thoracic SPNs located multiple segments rostral to the stimulation site. As noted in Methods, we targeted SPNs in T6-T8 and used the intact-cord preparation with an obliquely cut surface having a targeted caudal edge at T8. Retrogradely labelled SPNs were visualized and patched (n=20). Optical stimulation over the ventral surface of L1-L2 spinal cord elicited EPSPs (n=4), APs (n=5), or failed to elicit a response (n=11) in thoracic SPNs (typical example shown in Fig. 7). Backfilled SPNs had a similar rheobase value of 25 ± 10 pA as the presumed SPNs noted above, recorded in slice. Interestingly, in the n=30 SPNs from either slice or intact cord preparations, we noted a relationship between the recorded SPN’s ability to generate repetitive firing in response to current injection and whether optical stimulation elicited an excitatory response. Specifically, of the 15 SPNs that did not produce repetitive firing in response to a 1-s depolarizing current pulse, only 1 showed an excitatory response to optical stimulation. In contrast, 14/15 SPNs that demonstrated repetitive firing in response to current injection also showed an excitatory response to optical stimulation.

**Figure 7.**
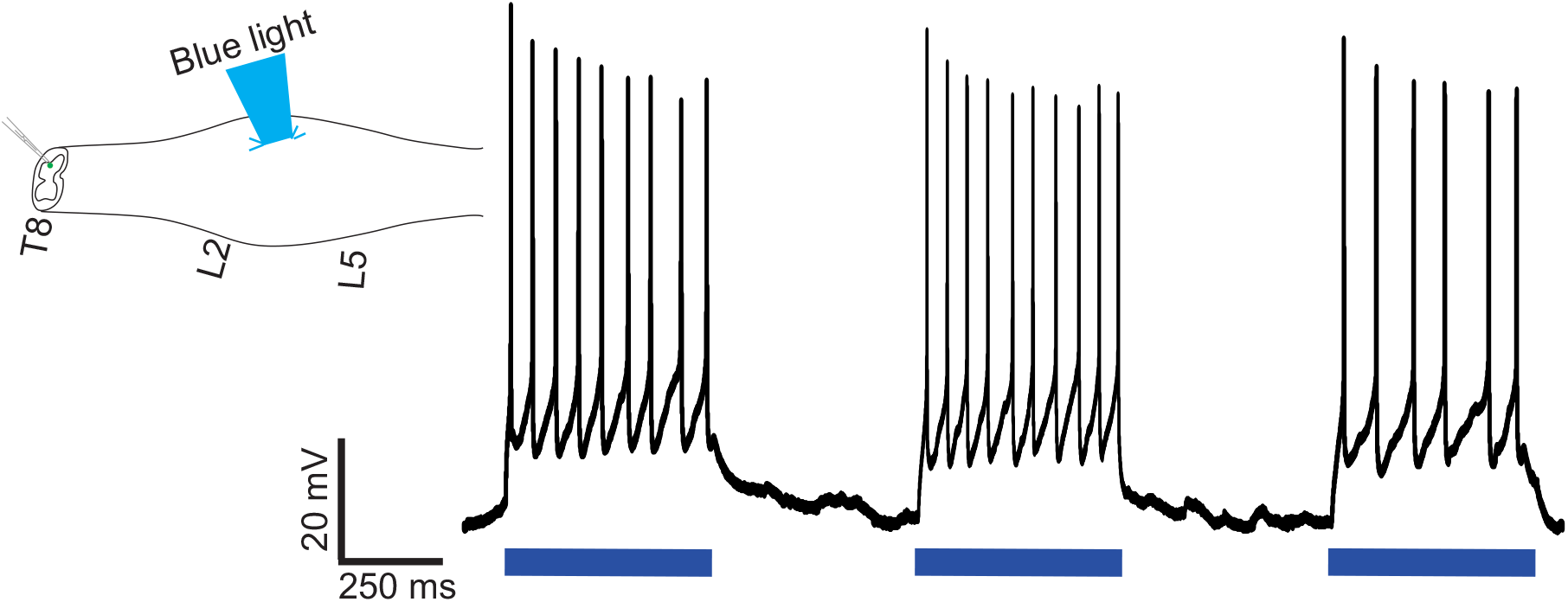
Optical stimulation of lumbar V3 neurons elicits action potentials in thoracic SPNs. In intact spinal cord preparations, thoracic SPNs labelled with RDA were visualized under fluorescence and whole cell patch clamp recordings were collected in response to optical stimulation of the ventral surface of the L2 spinal segment. In this example, lumbar V3 stimulation consistently evoked action potentials in the patched SPN. Representative example from one of the n =20 cells recorded from 15 mice of either sex.

Taken together, these anatomical and electrophysiological experiments provide compelling evidence of an ascending intraspinal excitatory neuronal projection from lumbar locomotor-related V3 neurons to thoracic sympathetic preganglionic neurons.

## DISCUSSION

Here, we demonstrate that locomotor related V3 INs project to and excite SPNs throughout the entire thoracic spinal cord and account for ∼20% of glutamatergic input to SPNs. We further demonstrate strong ascending input from lumbar V3 INs to thoracic SPNs (∼ 10% of the VGLut2 input), and, when optically stimulated, evokes post synaptic excitatory responses in SPNs. That is, we demonstrated a functional excitatory intraspinal connection between lumbar V3 INs and thoracic SPNs. We suggest that this newly described intraspinal connection provides an intraspinal capability to integrate locomotor and sympathetic function, which supports the appropriate activation of metabolic and homeostatic support during movement and exercise.

At the supraspinal level, clear evidence demonstrates the brainstem is capable of coordinating and integrating circuitry for movement with related needed sympathetic autonomic functions. Stimulation of the mesencephalic locomotor region (MLR) or subthalamic locomotor region not only results in locomotion but also increases respiratory function (Eldridge, Millhorn et al. 1981) and cardiovascular responses, in an activity and intensity dependent manner (Bedford, Loi et al. 1992).

Both of these increased sympathetic responses persisted when neuromuscular blockers were applied, suggesting a central source regulating both locomotor and autonomic functions (Chong and Bedford 1997). Even thinking about moving induces increases in BP and HR in humans temporarily paralyzed by curare, demonstrating similar mechanisms in human and animal models, and suggesting a critical link between motor and sympathetic autonomic systems at supraspinal levels (Gandevia, Killian et al. 1993). Confirming that these responses are mediated by a central source, two direct pathways from the MLR were recently identified, one projecting to glutamatergic neurons in the medulla that initiate movement, and one projecting to the cardio-respiratory centre located in the rostral ventral lateral medulla (RVLM) (Koba, Kumada et al. 2022). Further, opto- or chemogenetic activation of serotonergic neurons in the RVLM induce increases in systemic BP which precedes and outlasts bouts of simultaneously elicited locomotor activity, suggesting an integrative role for the brainstem in coordinating locomotor and autonomic support systems ((Armstrong, Nazzal et al. 2017), c.f. Fig. 4B in (Cowley 2018)).

Evidence for coordination within brainstem locomotor and autonomic circuitry has been demonstrated for glutamatergic Chx10 neurons located in the gigantocellular nucleus of the medulla. Anatomical and electrophysiological evidence indicates these neurons contribute to the generation and coordination of locomotion (Bretzner and Brownstone 2013, Bouvier, Caggiano et al. 2015, Schwenkgrub, Harrell et al. 2020, Usseglio, Gatier et al. 2020) and when ablated, respiration rate generated in the nearby pre-Botzinger complex is reduced (Crone, Viemari et al. 2012). Similarly, MLR stimulation increases blood pressure, heart rate and vasoconstriction prior to any rhythmic motor activity, even in the absence of movement in unanesthetized, decerebrate paralyzed rats (Bedford, Loi et al. 1992, Chong and Bedford 1997, Koba, Yoshida et al. 2005). Anatomical studies using pseudorabies virus injection into a variety of motor and sympathetic autonomic tissues demonstrated higher-order supraspinal neurons that function in both somato-motor and sympathetic systems, termed somato-motor-sympathetic neurons (Kerman, Enquist et al. 2003, Kerman, Akil et al. 2006, Kerman, Bernard et al. 2007, Kerman 2008). These findings suggest that communication and adequate activity in both brainstem autonomic and locomotor centres is sufficient and required for movement to occur. Based on evidence provided here, we propose a similar coordinating role between these systems within the spinal cord, and that V3 INs are involved in integrating and communicating between both autonomic SPNs and lumbar locomotor networks. Such integration between somatic motor and sympathetic autonomic systems appears to be a common feature at the afferent-spinal reflex level as well.

For example, Sato and colleagues described multiple somatic-motor and cutaneous non-nociceptive afferent evoked intra-segmental and multi-segmental sympathetic metabolic and homeostatic spinal reflexes in multiple species (Schmidt and Weller 1970, Sato 1973, Sato and Schmidt 1973, Sato 1987, Flett, Garcia et al. 2022). These reflexes are additional to the well described ‘pressor effect’ of lower limb afferents on BP regulation and include increases and decreases in catecholamine release from adrenal glands in anaesthetized preparations, dependent upon direction of mechanical stimulation of fur follicles (Sato and Schmidt 1973, Loewy and Spyer 1990, Schreihofer and Sved 2011), Similar to somatic motor reflexes, these sympathetic metabolic reflexes may become reversed or exaggerated after SCI due to a loss of descending control (reviewed in (Flett, Garcia et al. 2022)).

Our conceptual framework proposed that the spinal cord has the capability of integrating motor and sympathetic functions, and that this residual function may become critically important after SCI, when descending commands are removed (Cowley 2018, Flett, Garcia et al. 2022). Our recent scoping review supports this concept, as spinal electrical stimulation below injury targeted to lumbar regions to improve motor activity could also improve sympathetic autonomic functions, particularly in those with cervical level injury with their severely reduced or absent ability to activate sympathetic autonomic spinal neuronal circuitry (Flett, Garcia et al. 2022). We hypothesized the mechanisms mediating improved locomotor and autonomic function involved ascending propriospinal neural networks.

Ascending propriospinal neurons arising from the lumbar spinal cord can coordinate and entrain the locomotor rhythm on both thoracic respiratory neurons (Le Gal, Juvin et al. 2016) and the cervical CPG network (Juvin, Le Gal et al. 2012, Zhang, Shevtsova et al. 2022). Although we examined multiple glutamatergic neurons (V2a, V0D, V0V & V0C), including several that are classified as propriospinal, our results indicate that it may be V3 INs that provide a major source of spinal glutamatergic input to SPNs. V3 INs are ideal candidates to communicate and integrate locomotor and sympathetic functions as they have long ascending projections from lumbar to cervical spinal cord, and are involved in stability of locomotion and coordination between left and right hindlimbs and between hindlimbs and forelimbs (Zhang, Narayan et al. 2008, Blacklaws, Deska-Gauthier et al. 2015, Danner, Zhang et al. 2019, Zhang, Shevtsova et al. 2022), all of which is needed to sustain overground locomotion, particularly at higher intensities or durations.

We demonstrated that ∼20% of all VGluT2^+^ synaptic contacts on SPNs arise from V3 INs. As SPNs primarily receive glutamatergic input from neurons using the glutamatergic transporter VGluT2 (Llewellyn-Smith, Martin et al. 2007), this indicates that ∼20% of all glutamatergic inputs to SPNs are from V3 INs. Thus a large portion of excitatory input onto SPNs arises from V3 INs and about half this input appears to originate from lumbar V3 INs. Although it is known that SPNs receive intraspinal synaptic input from neurons using a variety of neurotransmitters including glutamate, GABA, and acetylcholine, little is known of the source of the neurons providing input to SPNs (Llewellyn-Smith 2009), and our findings represent an important first step in this identification process. It is likely that intraspinal input to SPNs also includes locally projecting spinal interneurons located in laminae V, VII and X (Llewellyn-Smith 2009). In our preliminary investigations, we did not observe any input from locally projecting Chx10^+^ spinal INs or from DBX^+^ spinal INs apposing SPNs. Thus, in addition to the ∼20% of glutamatergic input to SPNs from V3 INs, we conjecture that a large portion of remaining glutamatergic excitatory input to SPNs arise from brainstem centres including the RVLM.

Although we demonstrated that V3 INs provide a substantial portion of excitatory amino acid input to SPNs, we have not identified the final target tissue V3 IN input to SPNs activates. Like somatic motor systems, the autonomic input to bodily tissues typically follows a somatotopic organization, although an intact connection to T1 is required to activate a variety of sympathetic autonomic tissues including the heart, sweat glands and vasculature smooth muscle (Cowley 2018, Flett, Garcia et al. 2022). For instance, the greatest density of nerve terminals and fibres in the IML arising from Enkephalin expressing neurons occurs in T1-T8. For substance P expressing neurons the greatest density occurs between C8-T12 and T11-L1, and for serotonergic input, T1-T5 in cat (Krukoff, Ciriello et al. 1985). A similar distribution of serotonergic fibres within the IML was also seen in the rabbit, with the greatest density in T3-T6 and L3-L4 and minimal numbers T1 and T10-T12 (Jensen, Llewellyn-Smith et al. 1995). This differential rostro-caudal neuromodulatory input to SPNs, could permit specific functional groups of SPNs to respond in different ways to the same homeostatic challenge. Future experiments should examine the relationship between intraspinal projections and ultimate body targets of SPN output. We know, for example, that rostral thoracic SPNs via Superior Cervical Ganglia and Stellate Ganglion are involved in the regulation of upper body targets such as pupils, salivary glands, sweat glands of the head and arms, and the heart. SPNs in mid thoracic spinal cord via Celiac ganglion control mesenteric vasculature, gut motility and secretion and the adrenal medullae. SPNs in the caudal SPN aid in regulation of lower body organs such as urinary bladder and reproductive organs. In addition, there is a general somatotopic organization to sympathetic innervation of sweat glands, vascular smooth muscle and white adipose tissues that typically follows the dermatomes (below T1) (Cowley 2018).

Although preliminary, we observed a peak density of V3 IN contacts within T4-T7 segments, which via post-ganglionic sympathetic neurons innervate the heart and adrenal medulla. In addition, we also demonstrated that V3 INs innervate thoracic somatic MNs. Taken together, it is conceivable that in response to increased motor function, lumbar V3 INs provide additional excitation to SPNs to increase heart rate and stroke volume, increase release of circulating catecholamines from adrenal glands and fatty acid release from white adipose tissue, as well as activate thoracic MNs to meet the increased metabolic demands for respiration and posture during overground locomotion. In addition, one would expect less innervation of SPNs involved in regulating gut motility or reproductive organs as these are not vital for sustained movements or exercise. That is, there is likely a selective innervation of SPNs from neurons involved in locomotion, to ensure adequate organ and tissue support for movements.

Overall, this is the first demonstration of an anatomical and functional connectivity within the spinal cord between neurons involved in locomotion with neurons involved in sympathetic functions. We propose that similar to the brainstem, communication between locomotor and autonomic centres also occurs within the spinal cord. Long projecting ascending V3 propriospinal neurons may be a key neuron population ensuring that adequate metabolic and homeostatic resources are maintained during sustained rhythmic activity such as exercise. And that these neurons may be of therapeutic importance for improving autonomic and motor function after SCI.

## ACKNOWLEDGEMENTS

We thank Shannon Deschamps for assistance with surgical and post-operative animal care and with immunohistochemistry.

## AUTHOR CONTRIBUTIONS

KCC & JWC conceived the research project; CC, KCC & JWC conceived and designed experiments; CC, CVN, NS & JWC performed experiments; CC & NS analyzed data; CC, KCC & JWC interpreted results; CC, JWC prepared figures; CC, CVN, JWC and KCC drafted the manuscript; and JWC & KCC edited and revised manuscript. All authors approved the final version of the manuscript.

## LEGENDS

**Abstract Figure.**
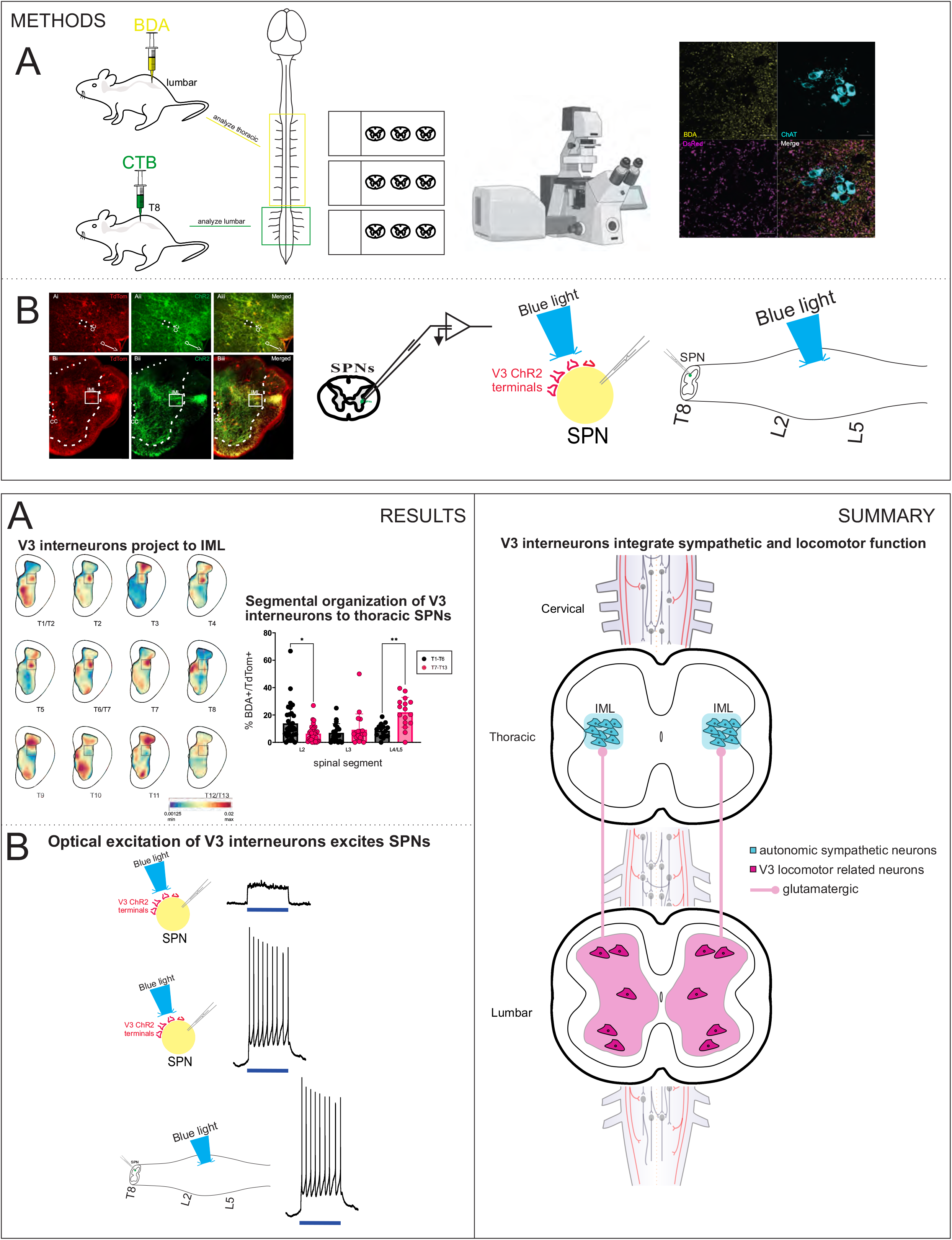
V3 locomotor-related neurons provide anatomical and functional input onto autonomic sympathetic neurons. A) Neuroanatomical tracing and immunohistochemistry methods schematic. B) Schematic methodology of electrophysiology protocols. C) Summary of results demonstrating that lumbar V3 interneurons provide ascending glutamatergic connections onto autonomic sympathetic neurons situated in the intermediate lamina (IML) of the thoracic spinal cord.

## REFERENCES

Adler, E. S., J. H. Hollis, I. J. Clarke, D. R. Grattan and B. J. Oldfield (2012). “Neurochemical characterization and sexual dimorphism of projections from the brain to abdominal and subcutaneous white adipose tissue in the rat.” J Neurosci 32(45): 15913–15921.

Al-Mosawie, A., J. M. Wilson and R. M. Brownstone (2007). “Heterogeneity of V2-derived interneurons in the adult mouse spinal cord.” Eur J Neurosci 26(11): 3003–3015.

Armstrong, K. E., M. Nazzal, X. Chen, K. Stecina and L. M. Jordan (2017). Chemogenic activation of parapyramidal brainstem neurons to evaluate motor consequences. Society for Neuroscience Abstracts, Washington DC, USA., Society for Neuroscience.

Bartness, T. J., Y. Liu, Y. B. Shrestha and V. Ryu (2014). “Neural innervation of white adipose tissue and the control of lipolysis.” Front Neuroendocrinol 35(4): 473–493.

Bartness, T. J., Y. B. Shrestha, C. H. Vaughan, G. J. Schwartz and C. K. Song (2010). “Sensory and sympathetic nervous system control of white adipose tissue lipolysis.” Mol Cell Endocrinol 318(1-2): 34–43.

Bartness, T. J., C. H. Vaughan and C. K. Song (2010). “Sympathetic and sensory innervation of brown adipose tissue.” Int J Obes (Lond) 34 Suppl 1: S36–42.

Bedford, T. G., P. K. Loi and C. C. Crandall (1992). “A model of dynamic exercise: the decerebrate rat locomotor preparation.” J Appl Physiol (1985) 72(1): 121–127.

Blacklaws, J., D. Deska-Gauthier, C. T. Jones, Y. L. Petracca, M. Liu, H. Zhang, J. P. Fawcett, J. C. Glover, G. M. Lanuza and Y. Zhang (2015). “Sim1 is required for the migration and axonal projections of V3 interneurons in the developing mouse spinal cord.” Dev Neurobiol 75(9): 1003–1017.

Blessing, W. W., Y. H. Yu and E. Nalivaiko (1999). “Raphe pallidus and parapyramidal neurons regulate ear pinna vascular conductance in the rabbit.” Neurosci Lett 270(1): 33–36.

Borowska, J., C. T. Jones, H. Zhang, J. Blacklaws, M. Goulding and Y. Zhang (2013). “Functional subpopulations of V3 interneurons in the mature mouse spinal cord.” J Neurosci 33(47): 18553–18565.

Bouvier, J., V. Caggiano, R. Leiras, V. Caldeira, C. Bellardita, K. Balueva, A. Fuchs and O. Kiehn (2015). “Descending Command Neurons in the Brainstem that Halt Locomotion.” Cell 163(5): 1191–1203.

Bretzner, F. and R. M. Brownstone (2013). “Lhx3-Chx10 reticulospinal neurons in locomotor circuits.” J Neurosci 33(37): 14681–14692.

Chacon, C. (2022). Sympathetic nervous system and locomotor integration from V3 spinal neurons. Master of Science Research, University of Manitoba.

Chong, R. K. and T. G. Bedford (1997). “Heart rate, blood pressure, and running speed responses to mesencephalic locomotor region stimulation in anesthetized rats.” Pflugers Arch 434(3): 280–284.

Chopek, J. W., F. Nascimento, M. Beato, R. M. Brownstone and Y. Zhang (2018). “Sub-populations of Spinal V3 Interneurons Form Focal Modules of Layered Pre-motor Microcircuits.” Cell Rep 25(1): 146–156 e143.

Chopek, J. W., F. Nascimento, M. Beato, R. M. Brownstone and Y. Zhang (2018). “Sub-populations of Spinal V3 Interneurons Form Focal Modules of Layered Pre-motor Microcircuits.” Cell Rep 25(1): 146–156.e143.

Chopek, J. W., Y. Zhang and R. M. Brownstone (2021). “Intrinsic brainstem circuits comprised of Chx10-expressing neurons contribute to reticulospinal output in mice.” J Neurophysiol 126(6): 1978–1990.

Conte, W. L., H. Kamishina and R. L. Reep (2009). “Multiple neuroanatomical tract-tracing using fluorescent Alexa Fluor conjugates of cholera toxin subunit B in rats.” Nat Protoc 4(8): 1157–1166.

Cowley, K. C. (2018). “A new conceptual framework for the integrated neural control of locomotor and sympathetic function: implications for exercise after spinal cord injury.” Appl Physiol Nutr Metab 43(11): 1140–1150.

Cowley, K. C., E. Zaporozhets and B. J. Schmidt (2008). “Propriospinal neurons are sufficient for bulbospinal transmission of the locomotor command signal in the neonatal rat spinal cord.” J Physiol 586(6): 1623–1635.

Cowley, K. C., E. Zaporozhets and B. J. Schmidt (2010). “Propriospinal transmission of the locomotor command signal in the neonatal rat.” Ann N Y Acad Sci 1198: 42–53.

Crone, S. A., K. A. Quinlan, L. Zagoraiou, S. Droho, C. E. Restrepo, L. Lundfald, T. Endo, J. Setlak, T. M. Jessell, O. Kiehn and K. Sharma (2008). “Genetic Ablation of V2a Ipsilateral Interneurons Disrupts Left-Right Locomotor Coordination in Mammalian Spinal Cord.” Neuron 60(1): 70–83.

Crone, S. A., J. C. Viemari, S. Droho, A. Mrejeru, J. M. Ramirez and K. Sharma (2012). “Irregular Breathing in Mice following Genetic Ablation of V2a Neurons.” J Neurosci 32(23): 7895–7906.

Crone, S. A., G. Zhong, R. Harris-Warrick and K. Sharma (2009). “In Mice Lacking V2a Interneurons, Gait Depends on Speed of Locomotion.” Journal of Neuroscience 29(21): 7098–7109.

Danner, S. M., N. A. Shevtsova, A. Frigon and I. A. Rybak (2017). “Computational modeling of spinal circuits controlling limb coordination and gaits in quadrupeds.” Elife 6.

Danner, S. M., S. D. Wilshin, N. A. Shevtsova and I. A. Rybak (2016). “Central control of interlimb coordination and speed-dependent gait expression in quadrupeds.” J Physiol 594(23): 6947–6967.

Danner, S. M., H. Zhang, N. A. Shevtsova, J. Borowska-Fielding, D. Deska-Gauthier, I. A. Rybak and Y. Zhang (2019). “Spinal V3 Interneurons and Left-Right Coordination in Mammalian Locomotion.” Front Cell Neurosci 13: 516.

Eldridge, F. L., D. E. Millhorn and T. G. Waldrop (1981). “Exercise hyperpnea and locomotion: parallel activation from the hypothalamus.” Science 211(4484): 844–846.

Flett, S., J. Garcia and K. C. Cowley (2022). “Spinal electrical stimulation to improve sympathetic autonomic functions needed for movement and exercise after spinal cord injury: a scoping clinical review.” J Neurophysiol 128(3): 649–670.

Gandevia, S. C., K. Killian, D. K. McKenzie, M. Crawford, G. M. Allen, R. B. Gorman and J. P. Hales (1993). “Respiratory sensations, cardiovascular control, kinaesthesia and transcranial stimulation during paralysis in humans.” J Physiol 470: 85–107.

Jensen, I., I. J. Llewellyn-Smith, P. Pilowsky, J. B. Minson and J. Chalmers (1995). “Serotonin inputs to rabbit sympathetic preganglionic neurons projecting to the superior cervical ganglion or adrenal medulla.” J Comp Neurol 353(3): 427–438.

Jessell, T. M. (2000). “Neuronal specification in the spinal cord: inductive signals and transcriptional codes.” Nat Rev Genet 1(1): 20–29.

Jordan, L. M., J. Liu, P. B. Hedlund, T. Akay and K. G. Pearson (2008). “Descending command systems for the initiation of locomotion in mammals.” Brain Res Rev 57(1): 183–191.

Juvin, L., J. P. Le Gal, J. Simmers and D. Morin (2012). “Cervicolumbar coordination in mammalian quadrupedal locomotion: role of spinal thoracic circuitry and limb sensory inputs.” J Neurosci 32(3): 953–965.

Kerman, I. A. (2008). “Organization of brain somatomotor-sympathetic circuits.” Exp Brain Res 187(1): 1–16.

Kerman, I. A., H. Akil and S. J. Watson (2006). “Rostral elements of sympatho-motor circuitry: a virally mediated transsynaptic tracing study.” J Neurosci 26(13): 3423–3433.

Kerman, I. A., R. Bernard, D. Rosenthal, J. Beals, H. Akil and S. J. Watson (2007). “Distinct populations of presympathetic-premotor neurons express orexin or melanin-concentrating hormone in the rat lateral hypothalamus.” J Comp Neurol 505(5): 586–601.

Kerman, I. A., L. W. Enquist, S. J. Watson and B. J. Yates (2003). “Brainstem substrates of sympatho-motor circuitry identified using trans-synaptic tracing with pseudorabies virus recombinants.” J Neurosci 23(11): 4657–4666.

Koba, S., N. Kumada, E. Narai, N. Kataoka, K. Nakamura and T. Watanabe (2022). “A brainstem monosynaptic excitatory pathway that drives locomotor activities and sympathetic cardiovascular responses.” Nature Communications 13(1): 5079.

Koba, S., T. Yoshida and N. Hayashi (2005). “Sympathetically induced renal vasoconstriction during stimulation of mesencephalic locomotor region in rats.” Auton Neurosci 121(1-2): 40–46.

Krukoff, T. L., J. Ciriello and F. R. Calaresu (1985). “Segmental distribution of peptide- and 5HT-like immunoreactivity in nerve terminals and fibers of the thoracolumbar sympathetic nuclei of the cat.” The Journal of comparative neurology. 240(1): 103–116.

Lanuza, G. M., S. Gosgnach, A. Pierani, T. M. Jessell and M. Goulding (2004). “Genetic identification of spinal interneurons that coordinate left-right locomotor activity necessary for walking movements.” Neuron 42(3): 375–386.

Le Gal, J. P., L. Juvin, L. Cardoit and D. Morin (2016). “Bimodal Respiratory-Locomotor Neurons in the Neonatal Rat Spinal Cord.” J Neurosci 36(3): 926–937.

Llewellyn-Smith, I. J. (2009). “Anatomy of synaptic circuits controlling the activity of sympathetic preganglionic neurons.” J Chem Neuroanat 38(3): 231–239.

Llewellyn-Smith, I. J., C. L. Martin, N. M. Fenwick, S. E. Dicarlo, H. L. Lujan and A. M. Schreihofer (2007). “VGLUT1 and VGLUT2 innervation in autonomic regions of intact and transected rat spinal cord.” J Comp Neurol 503(6): 741–767.

Llewellyn-Smith, I. J., K. D. Phend, J. B. Minson, P. M. Pilowsky and J. P. Chalmers (1992). “Glutamate-immunoreactive synapses on retrogradely-labelled sympathetic preganglionic neurons in rat thoracic spinal cord.” Brain Res 581(1): 67–80.

Llewellyn-Smith, I. J. and A. J. M. Verberne (2011). Central regulation of autonomic functions. New York, Oxford University Press.

Loewy, A. D. and K. M. Spyer (1990). Central Regulation of Autonomic Functions. Toronto, Ontario, Oxford University Press.

Lundfald, L., C. E. Restrepo, S. J. Butt, C. Y. Peng, S. Droho, T. Endo, H. U. Zeilhofer, K. Sharma and O. Kiehn (2007). “Phenotype of V2-derived interneurons and their relationship to the axon guidance molecule EphA4 in the developing mouse spinal cord.” Eur J Neurosci 26(11): 2989–3002.

Moran-Rivard, L., T. Kagawa, H. Saueressig, M. K. Gross, J. Burrill and M. Goulding (2001). “Evx1 is a postmitotic determinant of v0 interneuron identity in the spinal cord.” Neuron 29(2): 385–399.

Morrison, S. F., A. F. Sved and A. M. Passerin (1999). “GABA-mediated inhibition of raphe pallidus neurons regulates sympathetic outflow to brown adipose tissue.” Am J Physiol 276(2 Pt 2): R290–297.

Nwachukwu, C. V., C. Chacon, N. Shahsavani, J. W. Chopek and K. C. Cowley (2022). “Lumbar locomotor-related V3 interneurons project directly onto, and excite thoracic sympathetic preganglionic neurons.” Society for Neuroscience.

Pierani, A., L. Moran-Rivard, M. J. Sunshine, D. R. Littman, M. Goulding and T. M. Jessell (2001). “Control of interneuron fate in the developing spinal cord by the progenitor homeodomain protein Dbx1.” Neuron 29(2): 367–384.

Rybak, I. A., K. J. Dougherty and N. A. Shevtsova (2015). “Organization of the Mammalian Locomotor CPG: Review of Computational Model and Circuit Architectures Based on Genetically Identified Spinal Interneurons(1,2,3).” eNeuro 2(5).

Sato, A. (1973). “Spinal and medullary reflex components of the somatosympathetic reflex discharges evoked by stimulation of the group IV somatic afferents.” Brain Res 51: 307–318.

Sato, A. (1987). “Neural mechanisms of somatic sensory regulation of catecholamine secretion from the adrenal gland.” Adv Biophys 23: 39–80.

Sato, A. and R. F. Schmidt (1973). “Somatosympathetic reflexes: afferent fibers, central pathways, discharge characteristics.” Physiol Rev 53(4): 916–947.

Schmidt, R. F. and E. Weller (1970). “Reflex activity in the cervical and lumbar sympathetic trunk induced by unmyelinated somatic afferents.” Brain Res 24(2): 207–218.

Schreihofer, A. M. and A. F. Sved (2011). The ventrolateral medulla and sympathetic regulation of arterial pressure. Central regulation of autonomic functions. I. Llewellyn-Smith and A. J. Verberne. New York, Oxford University Press: xv, 401 p.

Schwenkgrub, J., E. R. Harrell, B. Bathellier and J. Bouvier (2020). “Deep imaging in the brainstem reveals functional heterogeneity in V2a neurons controlling locomotion.” Sci Adv 6(49).

Smith, J. E., A. S. Jansen, M. P. Gilbey and A. D. Loewy (1998). “CNS cell groups projecting to sympathetic outflow of tail artery: neural circuits involved in heat loss in the rat.” Brain Res 786(1-2): 153–164.

Strack, A. M., W. B. Sawyer, L. M. Marubio and A. D. Loewy (1988). “Spinal origin of sympathetic preganglionic neurons in the rat.” Brain Res 455(1): 187–191.

Szokol, K., J. C. Glover and M. C. Perreault (2008). “Differential origin of reticulospinal drive to motoneurons innervating trunk and hindlimb muscles in the mouse revealed by optical recording.” J Physiol 586(Pt 21): 5259–5276.

Talpalar, A. E., J. Bouvier, L. Borgius, G. Fortin, A. Pierani and O. Kiehn (2013). “Dual-mode operation of neuronal networks involved in left-right alternation.” Nature 500(7460): 85–88.

Usseglio, G., E. Gatier, A. Heuzé, C. Hérent and J. Bouvier (2020). “Control of Orienting Movements and Locomotion by Projection-Defined Subsets of Brainstem V2a Neurons.” Current Biology.

Zagoraiou, L., T. Akay, J. F. Martin, R. M. Brownstone, T. M. Jessell and G. B. Miles (2009). “A cluster of cholinergic premotor interneurons modulates mouse locomotor activity.” Neuron 64(5): 645–662.

Zhang, H., N. A. Shevtsova, D. Deska-Gauthier, C. Mackay, K. J. Dougherty, S. M. Danner, Y. Zhang and I. A. Rybak (2022). “The role of V3 neurons in speed-dependent interlimb coordination during locomotion in mice.” Elife 11.

Zhang, Y., S. Narayan, E. Geiman, G. M. Lanuza, T. Velasquez, B. Shanks, T. Akay, J. Dyck, K. Pearson, S. Gosgnach, C. M. Fan and M. Goulding (2008). “V3 spinal neurons establish a robust and balanced locomotor rhythm during walking.” Neuron 60(1): 84–96.

Ziskind-Conhaim, L. and S. Hochman (2017). “Diversity of molecularly defined spinal interneurons engaged in mammalian locomotor pattern generation.” J Neurophysiol 118(6): 2956–2974.

